# Reference-free microsatellite instability detection from tumor sequencing using intrasample variability modeling

**DOI:** 10.64898/2026.01.13.699258

**Authors:** Georgios Vlachos, Tina Moser, Mitesh Patel, James R. White, Ellen Heitzer, Luis A. Diaz

## Abstract

Microsatellite instability (MSI) is a key predictive biomarker across multiple tumor types, but current next-generation sequencing (NGS)-based callers oftendependon matched normals, large reference panels, or pretrained machine-learning models, which limits portability across assays and sequencing centers. We developed PROMIS (PROfiling of Microsatellite InStability), a tumor-only, reference-free pipeline that detects MSI by leveraging intrasample variability at predefined microsatellite loci. PROMIS models repeat-length distributions with a discrete mixture framework to distinguish stable germline configurations from unstable loci with additional allele populations. Locus-level classifications are aggregated into a continuous MSI score, defined as the fraction of unstable loci, which can then be used to classify samples as MSI-high or microsatellite-stable.

We benchmarked PROMIS in colorectal (CRC), endometrial (UCEC), and gastric (STAD) cancers from The Cancer Genome Atlas (TCGA). PROMIS achieved an overall AUC of 0.995 and cohort-specific AUCs of 1.00 in CRC and STAD and 0.999 in UCEC, comparable to established tools despite not using matched normals or pretrained models. Subsampling and in silico dilutions showed robust performance with substantially fewer loci and down to 3% tumor fraction. Finally, in prostate and CRC cell-free DNA (cfDNA) cohorts, including Illumina TSO500 data and an 18-gene panel, PROMIS yielded assay-agnostic MSI scores concordant with orthogonal tissue- and panel-based classifications.

## INTRODUCTION

Microsatellite instability (MSI) results from the failure of DNA mismatch repair (MMR) mechanisms, leading to insertion-deletion mutations in short tandem repeats across the genome. Loss of MMR function, typically through mutations in *MLH1*, *MSH2*, *MSH6*, or *PMS2*, creates a hypermutated phenotype that drives neoantigen production and strong immune infiltration. This biology explains the remarkable response of MSI-high (MSI-H) or dMMR tumors to immune checkpoint inhibitors (ICIs), establishing MSI as a key predictive biomarker in oncology (1,2).

MSI-H occurs across multiple tumor types, most frequently in colorectal, endometrial, and gastric cancers, but also in a small fraction of other solid malignancies (3). Its pan-cancer relevance was underscored by the U.S. FDA’s tumor-agnostic approval of PD-1 blockade for MSI-H/dMMR cancers (4). Accurate and scalable detection of MSI status is therefore essential for both clinical decision-making and biomarker research.

Historically, MSI testing has relied on PCR fragment analysis of microsatellite loci and immunohistochemistry (IHC) for MMR proteins (5). While these methods are broadly concordant, they are labor-intensive, low-throughput, and provide limited genomic context. Next-generation sequencing (NGS) now enables genome-wide MSI assessment, supporting the development of computational tools that infer instability directly from sequencing data.

Early computational methods such as MSIsensor and MANTIS compared repeat-length distributions between tumor and matched-normal samples to identify unstable loci with high accuracy (6,7). However, these methods depend on matched normal DNA, which is often unavailable in clinical sequencing. Later tumor-only methods, including MSIsensor-pro (8), addressed this limitation by comparing tumor data to panels of normals or by modeling polymerase-slippage events statistically. More recent machine-learning and deep-learning approaches, such as MiMSI (9), further improved sensitivity, particularly in low-purity samples, but rely on pretrained models that may not generalize across sequencing panels or populations. Specialized tools like MSICare (10) achieve excellent accuracy in specific cancer types but are not broadly applicable.

Despite these advances, current MSI callers face three persistent challenges. They often rely on matched normal samples or large reference datasets, depend on pretrained models that limit interpretability and cross-platform portability, and reduce MSI to a binary label without readily revealing locus-specific patterns of instability that may hold biological or clinical relevance. A truly reference-free, tumor-only, and interpretable approach remains lacking.

To address this gap, we developed PROMIS (PROfiling of Microsatellite InStability), a novel bioinformatics pipeline that detects MSI directly from tumor sequencing data without matched normals, panels of normals, or machine-learning retraining. PROMIS leverages intrasample variability, treating each tumor as its own internal control. By modeling repeat-length distributions at predefined microsatellite loci using a discrete mixture modeling framework (11), PROMIS distinguishes stable monomorphic loci from unstable loci exhibiting multiple discrete allele populations. This statistical modeling accounts for natural polymorphism and sequencing noise, yielding a transparent, locus-level classification and a global MSI score based on the fraction of unstable loci per sample.

PROMIS is implemented as a fully automated and reproducible Snakemake workflow compatible with whole-genome, whole-exome, and targeted sequencing panels (12). Its modular design supports both tissue and circulating cell-free DNA (cfDNA) analyses, incorporating stringent read-level filtering, locus extraction, mixture-model fitting, and sample-level scoring. Each call is traceable and visualizable, ensuring interpretability and clinical auditability.

Benchmarking across The Cancer Genome Atlas (TCGA) cohorts of colorectal (CRC), endometrial (UCEC), and gastric cancers (STAD) demonstrated that PROMIS achieves sensitivity and specificity comparable to or exceeding state-of-the-art tools. PROMIS maintains high accuracy at low tumor fractions and extends MSI detection to cfDNA datasets, where traditional tools often fail.

Collectively, PROMIS represents a control-agnostic, sample-agnostic, and assay-agnostic solution for robust MSI detection. By combining transparent statistical modeling with full compatibility across sequencing modalities, PROMIS bridges the gap between research-grade MSI discovery and practical, scalable diagnostics.

## RESULTS

### Overview of the PROMIS pipeline

PROMIS is a tumor-only, reference-free bioinformatics pipeline for detecting MSI from DNA sequencing data. The workflow leverages intrasample variability, using each tumor as its own internal reference to identify unstable microsatellite loci.

PROMIS accepts aligned BAM files from whole-genome, whole-exome, or targeted-panel sequencing and executes four sequential stages **(Figure 1B)**. First, reads overlapping predefined microsatellite loci are extracted from the tumor BAM file. Second, stringent quality filters remove low-confidence reads and alignment artifacts. Third, repeat-length distributions at each locus are modeled using a discrete mixture model to detect multimodal allele populations indicative of instability. Finally, locus-level classifications are aggregated to compute a global MSI score representing the proportion of unstable loci within each sample.

**Figure 1:**
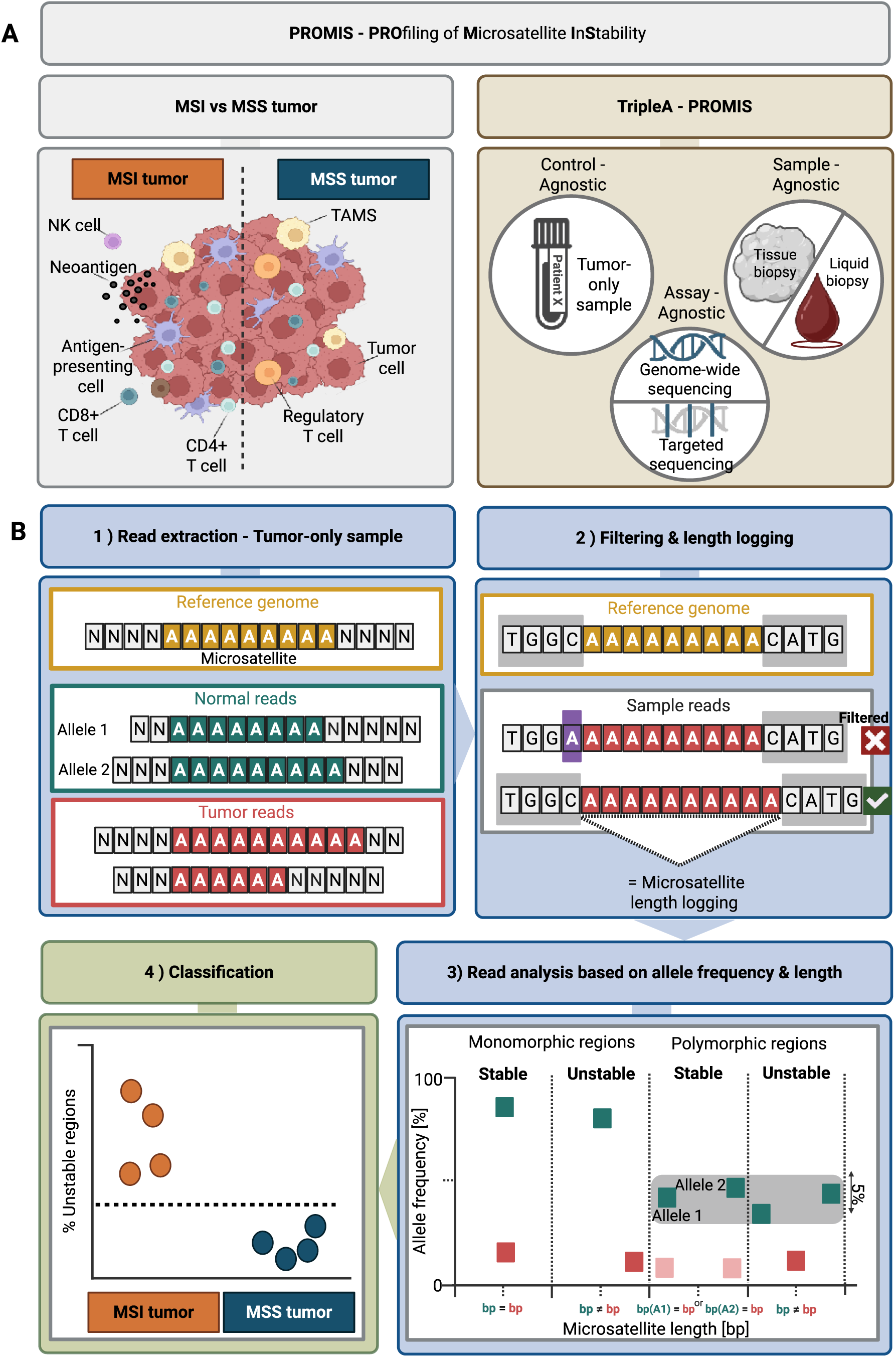
Overview of the PROMIS pipeline for tumor-only microsatellite instability (MSI) detection from next-generation sequencing data. PROMIS (PROfiling of Microsatellite InStability) is a control-agnostic, sample-agnostic, and assay-agnostic (“TripleA”) framework applicable to both tissue and liquid biopsy samples, and compatible with genome-wide (WES, WGS) or targeted sequencing panels. (A): Left: schematic representation of MSI vs MSS tumors, highlighting the immunogenic neoantigen landscape and tumor microenvironment. Right: the TripleA design principles ensure broad applicability across assays and sample types. ( B): PROMIS workflow steps. (1) Reads overlapping predefined microsatellite loci are extracted from tumor-only samples. (2) Reads undergo stringent quality filtering, where an “X” indicates a filtered-out read and a “✓” a retained read, followed by microsatellite length logging. (3) Allele frequency and length distributions are analyzed. The x-axis represents microsatellite length in base pairs (bp): bp = bp denotes equal lengths across alleles (stable), while bp ≠ bp denotes unequal lengths indicative of instability. Both monomorphic regions (normally uniform repeats) and polymorphic regions (with natural allelic variation) are modeled. A discrete mixture modeling approach distinguishes stable loci, exhibiting expected polymorphisms, from unstable loci with multiple distinct allele populations. (4) The proportion of unstable loci is calculated, enabling robust binary classification into microsatellite-stable (MSS) and MSI-high tumors.

PROMIS was benchmarked using 702 high-quality, well-annotated exonic microsatellite loci curated from the MOSAIC reference set (13), ensuring robust and consistent coverage across WES and WGS samples. The pipeline also includes a preprocessing module that automatically identifies which microsatellite loci are covered in each dataset based on the sequencing design and reference genome version. This step enables seamless adaptation of PROMIS to different panels or capture assays and ensures that only confidently represented loci are retained for downstream analysis. Each module produces interpretable outputs, including read-level statistics, per-locus instability calls, and visual summaries of allele-length distributions. The workflow is computationally efficient, completing a typical exome in under two minutes and scaling linearly with the number of loci analyzed. Multiple samples can be processed in parallel, allowing high-throughput execution for both large-scale research studies and clinical diagnostics.

### Performance of PROMIS across tumor types

We benchmarked PROMIS against established MSI callers using TCGA cohorts of CRC, UCEC, and STAD. PROMIS instability scores showed strong concordance, with Pearson correlation coefficients (r) of 0.90 for both comparisons (p < 10⁻¹⁰) **(Figure 2C)**. This high degree of correlation confirms that PROMIS quantitatively recapitulates the behavior of established MSI callers. When excluding *POLE* mutated samples, PROMIS also correlated strongly with tumor mutational burden and mutation count, reflecting the underlying MSI-associated hypermutation phenotype **(Supplementary Figure 1)**.

**Figure 2:**
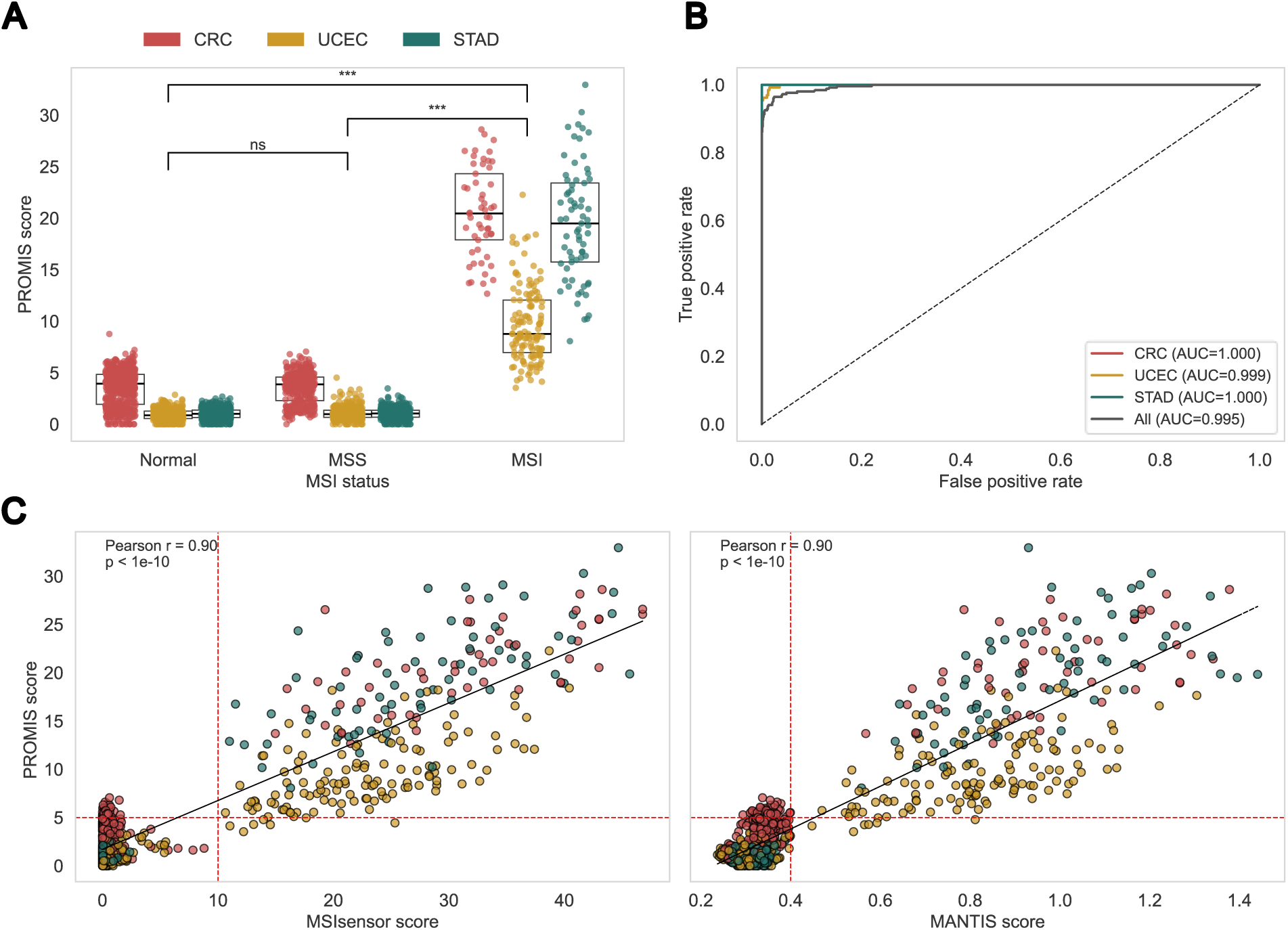
Performance and cross-platform validation of the PROMIS pipeline. **(A)** PROMIS scores by MSI status. Each point represents a tumor sample, stratified by consensus MSI status (MSS or MSI) defined by concordant MSIsensor and MANTIS calls. Points are colored by cancer type (CRC, UCEC, STAD). Only samples with classifier agreement and mean coverage >30× are included. Normal tissue scores are shown in the left. **(B)** Receiver operating characteristic (ROC) curves for PROMIS performance in different cancer cohorts. **(C)** Correlation of PROMIS scores with established MSI classifiers. Left: scatterplot of PROMIS vs. MSIsensor score; right: PROMIS vs. MANTIS score. Points are colored by cancer type. Red dashed lines indicate the classification thresholds for each method. Pearson correlation coefficients (r) and p-values are shown.

Across all tumor types, PROMIS accurately classified MSI status, achieving an overall area under the curve (AUC) of 0.99 **(Figure 2B)**. Using a representative instability threshold of 5% unstable loci, sensitivity ranged from 95% in endometrial to 100% in colorectal and gastric cancers, with an overall sensitivity of 97% across cohorts. Specificity was 100% for endometrial and gastric tumors and 87% for colorectal tumors, the latter reflecting higher background instability in MSS and normal colorectal tissues **(Figure 2A)**, a phenomenon previously reported in this cancer type (14). Increasing the CRC MSI classification threshold to 10% for colorectal tumors achieved 100% sensitivity and specificity.

We compared PROMIS performance with published AUC values from established MSI detection tools across TCGA and MSK-IMPACT cohorts **(Table 1)**. PROMIS achieved an overall AUC = 0.995, with cohort-specific values of 1.00 for CRC and STAD and 0.999 for UCEC. These results are comparable to MSIsensor-pro (AUC = 0.994), MANTIS (AUC = 0.986), and MSIsensor (AUC = 0.989), and consistent with the deep-learning model MiMSI (AUC = 0.972) (8,9). Together, these findings indicate that PROMIS delivers accuracy on par with current state-of-the-art approaches while remaining independent of matched-normal data or pretrained models.

**Table 1:**
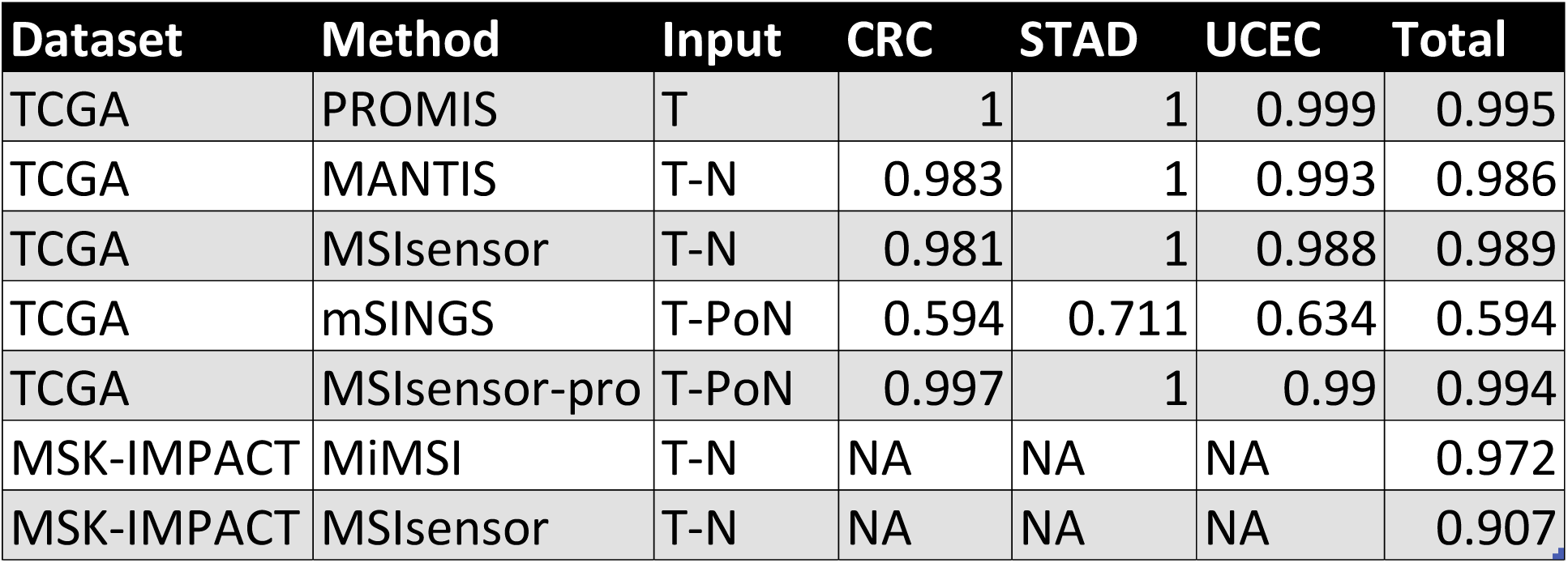
Comparative AUC performance of MSI detection methods across cancer types. Area under the ROC curve (AUC) values are shown for each method and dataset, stratified by cancer type: colorectal cancer (CRC), stomach adenocarcinoma (STAD), and uterine corpus endometrial carcinoma (UCEC), as well as the overall total. Methods are grouped by input requirements: tumor-only (T; PROMIS), matched tumor–normal pairs (T–N; MANTIS, MSIsensor, MiMSI, MSIsensor on MSK-IMPACT), and tumor plus panel of normals (T–PoN; mSINGS, MSIsensor-pro). NA: not available for that cohort.

### Robustness across genomic subsets and tumor purity

To evaluate the robustness of PROMIS across different sequencing contexts, we assessed its performance using variable subsets of microsatellite loci and simulated tumor purities. Subsampling analyses showed that classification accuracy remained stable even when the number of analyzed loci was reduced fivefold, with AUC values consistently above 0.93 **(Figure 3A)**. A moderate drop in AUC was observed for UCEC samples, indicating that a larger number of loci is required for optimal classification in this tumor type, consistent with the narrower separation between MSI and MSS cases compared with colorectal and gastric cancers. Moreover, the informative loci differed between UCEC and CRC/STAD, as reflected by their separation in two-dimensional principal component space **(Figure 3B)**. Overall, these findings indicate that PROMIS maintains reliable performance on targeted panels containing a limited number of microsatellites while capturing tumor-type specific instability patterns.

**Figure 3:**
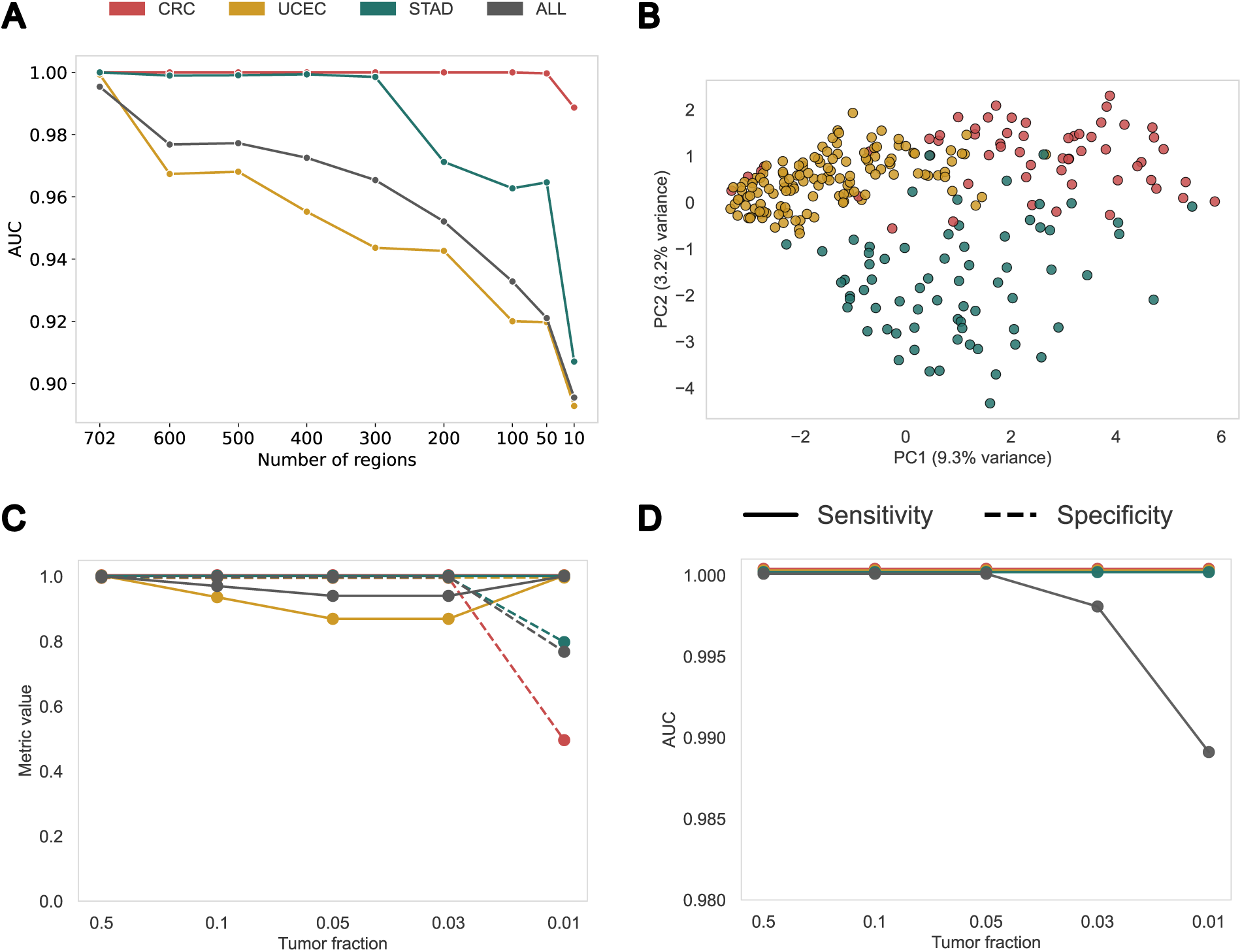
Robustness of PROMIS across genomic subsets and tumor fractions. **(A)** Area under the ROC curve (AUC) for PROMIS classification as a function of the number of microsatellite regions used. Line plots show CRC, UCEC, STAD, and pooled tumors (ALL). Only subsets with >25 regions are included. The x-axis is reversed to show decreasing region counts from left to right. **(B)** Principal component analysis (PCA) of instability profiles across recurrent microsatellite loci in consensus MSI-high tumors. Each point represents atumor sample, colored by cancer type (CRC, UCEC, STAD). Variance explained by each component is indicated on the axes. **(C)** PROMIS performance at decreasing tumor fractions. AUC values are shown for CRC, UCEC, STAD, and pooled tumors (ALL) across four dilution levels (50%, 10%, 5%, 3%, 1% tumor fraction). Only samples with concordant MSIsensor and MANTIS calls are included. A subset of 50 representative samples were used for dilution testing. **(D)** Sensitivity and specificity of PROMIS across decreasing tumor fractions. Solid and dashed lines represent sensitivity and specificity, respectively, for CRC, UCEC, STAD, and pooled tumors (ALL).

To assess sensitivity to tumor purity, we generated *in silico* dilution series for 50 representative cases spanning the full range of PROMIS scores **(Supplementary Figure 2)**, from clearly MSI to borderline and MSS **(Figure 3C–D)**. PROMIS retained near-perfect separation of MSI and MSS cases across all tumor types down to 3 % tumor fraction. Even at 1 %, PROMIS continued to produce measurable MSI signal, although classification accuracy decreased slightly. When applying the PROMIS decision threshold of 5, sensitivity remained high across dilutions, whereas specificity declined at the lowest tumor fractions, reflecting score inflation due to relaxed detection stringency. These findings confirm that PROMIS is highly robust across tumor purities and suitable for low-input sequencing contexts such as cfDNA or small clinical biopsies.

### Application to cfDNA and targeted panels

We next evaluated the performance of PROMIS in cfDNA sequencing data and targeted gene panels, which present additional challenges due to low tumor fraction and limited genomic coverage. PROMIS was applied to prostate cancer cfDNA samples analyzed across three assay types; whole-exome sequencing, whole-genome sequencing, and an amplicon panel targeting 18 genes. MSI status for these samples was determined using matched tumor-normal MSIsensor classifications from WES. PROMIS reliably identified the single MSI-positive case across all three assays, with instability fractions consistent with the reference calls, an observation in line with the low frequency of MSI reported in prostate cancer **(Figure 4A)**.

**Figure 4:**
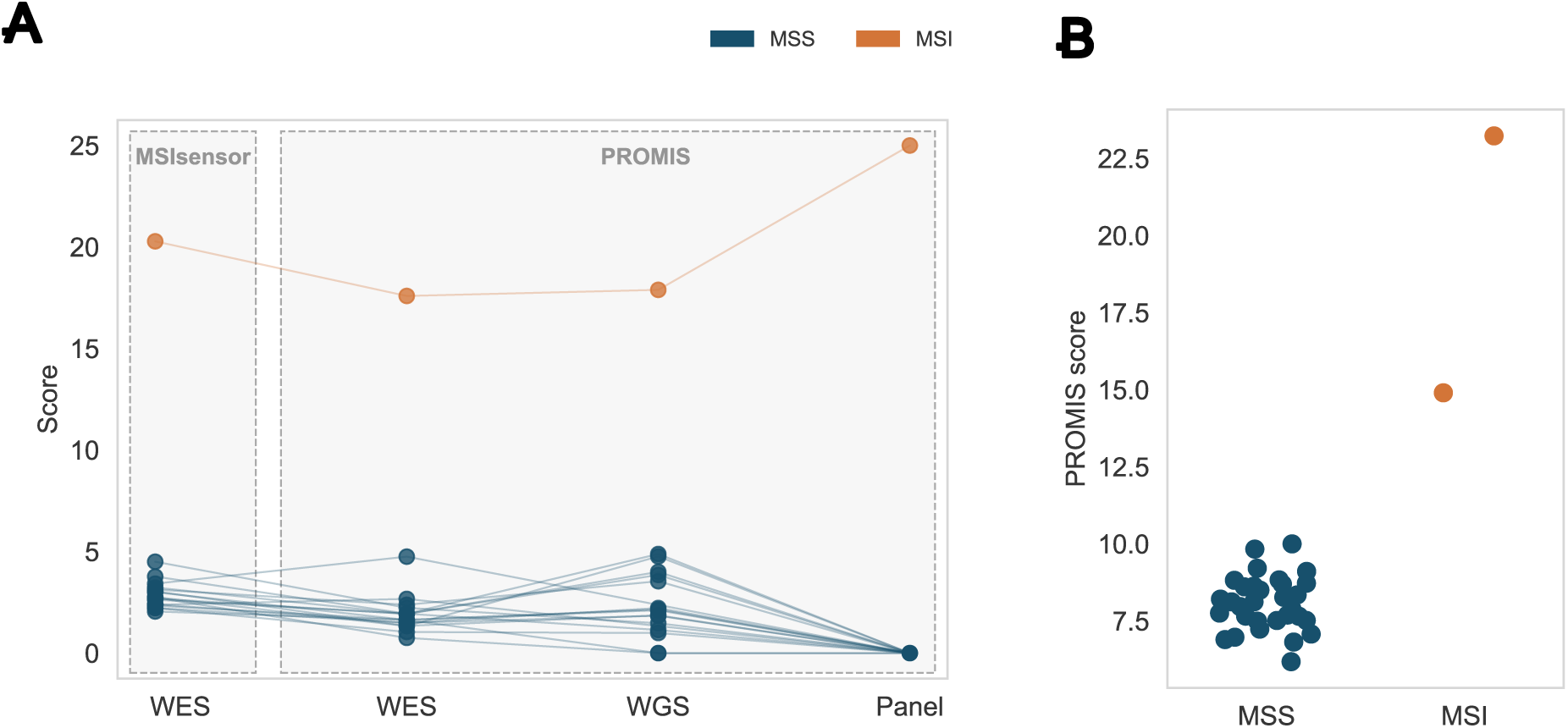
PROMIS performance in liquid biopsy assays. **(A)** Comparison of PROMIS scores with MSIsensor in prostate cancer cfDNA (WES and 20x WGS) and targeted panel (18 genes, QIAseq-HRD). Each point represents a patient sample, colored by consensus MSI status (MSS or MSI). Lines connect scores across methods for the same sample. Gray dashed boxes highlight MSIsensor (left) and PROMIS (right) groups of methods. **(B)** PROMIS scores in colorectal cancer cfDNA (TSO500 panel). Each point represents a patient sample, stratified by MSI status (MSS or MSI). Scores are expressed as the percentage of unstable microsatellite regions.

In addition, PROMIS was evaluated in CRC cfDNA samples profiled using the TSO500 targeted sequencing assay, with MSI status derived from the assay report. PROMIS scores clearly separated MSI from MSS cases **(Figure 4B)**. As previously noted, CRC samples displayed slightly higher baseline instability levels, indicating that a 10% threshold provides optimal discrimination for this tumor type. Together, these findings demonstrate that PROMIS performs consistently across cfDNA assays and supports accurate MSI assessment in liquid biopsy settings.

## DISCUSSION

We developed PROMIS, a reference-free, tumor-only pipeline for detecting MSI from NGS data. By modeling intrasample variation in microsatellite repeat lengths, PROMIS infers MSI status without matched normals, panels of normals, or pretrained models. The method demonstrated strong concordance with established tools and remained stable across sequencing contexts, including low-tumor-fraction and cfDNA samples. Together, these results show that PROMIS enables fast, interpretable, and assay-independent MSI assessment suitable for both research and clinical applications.

MSI detection has evolved from PCR- and IHC-based assays to computational analyses integrated within sequencing workflows. Established algorithms such as MSIsensor, MANTIS, and MSIsensor-pro perform well but rely on matched controls or fixed reference models, which can limit portability across datasets and platforms. Tumor-only deep-learning methods such as MiMSI improve sensitivity in low-purity data but require extensive model training, are prone to batch effects, and may need retraining for different sequencing centers or assay designs. Moreover, their decision boundaries are not easily interpretable. PROMIS provides a complementary alternative: a fully statistical framework that quantifies instability within each tumor sample rather than relative to external references. This design reduces dependencies, improves transparency, and enables consistent MSI calling across diverse sequencing modalities.

Although PROMIS performs consistently across tumor types, several limitations should be acknowledged. MSI status for TCGA samples was derived from the consensus of MSIsensor and MANTIS, with discrepant cases excluded from benchmarking. Examination of these excluded samples showed that those classified as MSI by MANTIS and borderline by MSIsensor were typically called MSI by PROMIS, whereas cases labeled borderline by MANTIS and MSS by MSIsensor were classified as MSS **(*Supplementary Figure 3*)**, suggesting that PROMIS generally produced the more concordant calls. Performance also depends on the number and diversity of microsatellite loci analyzed; UCEC showed slightly reduced separation between MSI and MSS groups, indicating that tumor-specific or expanded panels may further enhance sensitivity. The method assumes a moderate degree of normal DNA admixture or subclonality, as this internal heterogeneity provides the reference signal for stable allele lengths. In extremely high-purity samples (>95% tumor content), or in cell lines lacking allelic diversity, a fully homozygous unstable locus may be missed. Conversely, very low-purity samples (<1% tumor content) can fall below detection limits, where instability signals become indistinguishable from noise. These edge cases are uncommon in clinical sequencing but represent practical boundaries of the current implementation.

Future development of PROMIS will focus on expanding its flexibility and integration within broader genomic pipelines. The modular structure allows straightforward adaptation to new sequencing panels and capture designs, and the preprocessing module can be extended to dynamically adjust locus selection based on coverage or assay content. Importantly, by examining the characteristics of specific unstable loci, such as TA-repeat instability associated with WRN helicase dependency (15,16), PROMIS could extend beyond binary MSI classification to inform therapeutic stratification. This functionality can be readily implemented within the existing framework by annotating instability events to gene or motif contexts. Finally, we aim to enhance the pipeline’s accessibility and user-friendliness by incorporating Bioconda-based environment management and ensuring ready implementation in Nextflow workflows, while maintaining full open-source availability (17,18). These developments will facilitate broader community adoption, benchmarking, and collaborative improvement of PROMIS.

PROMIS provides a reliable and reproducible framework for detecting microsatellite instability directly from tumor sequencing data. By removing the need for matched normals or pretrained models, it simplifies MSI calling while maintaining statistical interpretability across platforms and sample types. Its scalability, modular design, and compatibility with cfDNA workflows make PROMIS a practical tool for integrating MSI detection into modern genomic pipelines.

## METHODS

### Data sources and MSI reference labels

WES data were obtained from TCGA (RRID:SCR_003193) for CRC (COAD/READ), UCEC, and STAD. Aligned and preprocessed BAM files (hg38) were downloaded from the Genomic Data Commons (GDC, RRID:SCR_014514). Samples with mean coverage below 30× were excluded from analysis.

Reference MSI classifications were derived from published TCGA consensus annotations generated using MSIsensor (RRID:SCR_006418) and MANTIS, applying their standard thresholds of 10 and 0.4, respectively (6,7). Only samples with concordant calls between both tools were used as benchmark reference labels. Discrepant cases were excluded from the validation set.

For cfDNA analyses, prostate cancer samples were analyzed using WES, WGS, and targeted amplicon panel sequencing. All datasets were aligned and processed according to the GATK Best Practices workflow (RRID:SCR_001876), including alignment with BWA-MEM2 (RRID:SCR_022192) (19), sorting, duplicate marking, and base quality recalibration, using the hg38 reference genome. MSI status for these samples was determined from matched tumor - normal MSIsensor calls on the WES data.

Colorectal cfDNA samples were analyzed using the Illumina TSO500 targeted sequencing assay. FASTQ files were processed on the Illumina DRAGEN Bio-IT platform, which performs alignment (hg19), variant calling, and MSI scoring within the TSO500 analysis suite. MSI status for these samples was obtained directly from the assay report.

### Workflow overview

PROMIS operates on aligned BAM files and a reference set of microsatellite loci defined in the corresponding genome build. The workflow proceeds through sequential stages, each focused on a specific analytical task. All parameters like base and mapping quality or MSI calling thresholds can be readily adjusted through a configuration file.

#### 1. Read extraction and filtering

Sequencing reads overlapping predefined microsatellite loci are identified using coordinate-based queries. Only reads that span the full repeat region with at least 4 bp of flanking sequence on each side are retained. Quality control filters exclude reads with mapping quality below 40 or mean base quality below 20 within the repeat tract. Reads containing soft-clipped alignments, supplementary mappings, or mismatched repeat motifs are also discarded to prevent artifacts from misalignments or sequencing errors. The retained reads are used to reconstruct the observed repeat length per locus, producing a read-level repeat-length profile for each sample.

#### 2. Repeat-length quantification

For each microsatellite locus, the distribution of observed repeat lengths across all high-quality reads is computed. Stable loci typically display one or two dominant alleles (corresponding to homozygous or heterozygous germline configurations), while unstable loci exhibit additional repeat-length variants reflecting somatic indel events. Loci with fewer than 50 total reads are flagged as low-confidence and excluded from downstream modeling.

#### 3. Instability modeling and locus classification

For each locus *l*, the likelihood of observing a repeat length *x* across *n* reads is expressed as:

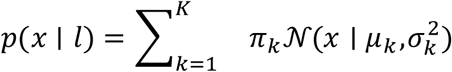

where:

- *x* represents an individual repeat length observation (in base pairs)
- *K* is the number of mixture components (ranging from 1 to 4), representing distinct allelic populations
- π_k_ is the mixing coefficient (relative proportion) of reads assigned to component *k*, with ∑π_k_ = 1 and 0 ≤ π_k_ ≤ 1
- μ_k_ is the mean repeat length of component *k* (in base pairs)
- (σ_k_)^2^ is the variance of component *k*, capturing the spread of repeat lengths around the mean
- 𝒩(x | μ_k_, (σ_k_)^2^) is the Gaussian probability density function for component *k*

Although repeat lengths are discrete integers, a continuous Gaussian approximation provides a stable estimate of distinct allelic modes under moderate coverage conditions.

Model parameters Θ = {π₁…π_K, μ₁…μ_K, σ₁²…σ_K²} were estimated using the Expectation-Maximization (EM) algorithm implemented in scikit-learn’s GaussianMixture class (RRID:SCR_002577). To ensure numerical stability and prevent singular covariance matrices in cases of limited variance, we applied regularization by adding a small constant to the diagonal of covariance matrices:

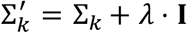

where:

- Σ_k_ is the covariance matrix for component *k*
- λ = 1×10⁻² is the regularization coefficient (reg_covar parameter)
- **I** is the identity matrix
- Σ_k’_ is the regularized covariance matrix used in model fitting

Each model was initialized 5 times (n_init_ = 5) with different random starting values, and the solution achieving the highest log-likelihood was retained to mitigate local optima in the EM algorithm.

The optimal number of mixture components *K* is selected by minimizing the Bayesian Information Criterion (BIC):

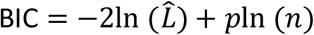

where:

- 𝐿^is the maximized likelihood of the fitted model, representing how well the model explains the observed data
- *p* is the number of free parameters in the model (for a GMM with *K* components: *p* = *K* - 1 mixing weights + *K* means + *K* variances = 3*K* - 1)
- *n* is the number of reads supporting the locus

To prevent overfitting and distinguish biologically meaningful germline heterozygosity from technical noise or somatic instability, a secondary allele ( μ₂) was retained only if all of the following criteria were satisfied:

1. Adequate representation: 𝜋_2_ ≥ 𝑓_min_ where f_min_ = 0.25 (minimum fraction of reads, default parameter)
2. Balanced allelic ratio: 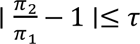 where τ = 0.05 (balance tolerance, default parameter)

These stringent criteria ensure that secondary components represent genuine heterozygous germline variants (expected ratio ≈ 1:1) rather than low-level somatic mutations or sequencing artifacts. A locus is classified as heterozygous when both alleles meet these thresholds; otherwise, it is treated as homozygous with a single stable length.

#### 4. Locus-level instability scoring and sample-level MSI computation

Following mixture model fitting, each microsatellite locus was evaluated for evidence of instability based on its repeat-length distribution. A locus was designated as unstable when a sufficient fraction of reads deviated from the inferred stable allelic modes by at least one repeat unit. Deviations were quantified using two complementary criteria: a minimum number of deviating reads and a minimum percentage of the total reads supporting an alternate allele length. By default, loci with fewer than 50 total reads were excluded from analysis to minimize stochastic effects at low coverage, and a locus was called unstable if at least five deviating reads or ≥1% of all reads supported adistinct allele length. All thresholds, including minimum read depth, minimum deviation counts, and percentage cut-offs, are user-configurable within the PROMIS configuration file.

For each sample, PROMIS aggregates locus-level classifications to compute a global instability score, defined as the fraction of evaluated loci classified as unstable. This continuous MSI score reflects the overall burden of microsatellite instability within the tumor and can be readily converted to a binary MSI-high or microsatellite-stable (MSS) status by applying a user-defined cut-off.

The pipeline outputs comprehensive summary tables containing per-locus statistics (lengths of deviating reads, number of deviating reads, instability flag) and sample-level metrics, allowing both automated classification and manual inspection of borderline cases.

#### 5. Microsatellite locus definition and preprocessing

PROMIS analyzes a curated set of 702 high-quality microsatellite loci derived from the MOSAIC reference dataset (13), encompassing mono-, di-, tri-, and tetranucleotide repeats across exonic and clinically relevant regions.

For analysis of new assays, PROMIS automatically determines which of these loci are covered by cross-referencing the sequencing regions or aligned reads with the reference genome. Loci with insufficient read depth or mapping quality are excluded from downstream modeling. This preprocessing step enables PROMIS to adapt dynamically to WGS, WES, and targeted panels without requiring retraining or recalibration, ensuring consistent analysis across different assay designs.

### Assessment of number of regions needed for reliable calls

To evaluate the impact of locus number on MSI classification accuracy, all microsatellite regions were ranked according to their discriminatory power, callability, and background stability. For each locus, ROC AUC values were computed using consensus MSI labels from MSIsensor, MANTIS, and TCGA clinical annotations, together with callability rates (coverage >30×) and instability frequency in MSS samples. A composite score integrating these metrics was bootstrapped 100 times to obtain mean stability ranks, and loci were sorted accordingly. Nested subsets containing the top 600, 500, 400, 300, 200, 100, 50 and 10 loci were then used to recompute PROMIS MSI scores for all samples.

### Simulation of tumor purity

To evaluate PROMIS performance at varying tumor purities, in silico dilution series were generated by mixing tumor and matched-normal sequencing reads at predefined fractions. For each tumor, synthetic BAM files were created at 50%, 10%, 5%, and 3% tumor f ractions while maintaining total read depth constant. Mixing was performed read-pair aware, preserving original pairing, read-group metadata, and sequencing quality.

A representative set of 50 samples was selected across tumor types and MSI status categories, including high-MSI, borderline-MSI, borderline-MSS, and MSS cases **(Supplementary Figure 2)**. Original tumor purity estimates were obtained from ABSOLUTE values (RRID:SCR_005198) (20), provided in the TCGA metadata.

For MSI classification, the standard parameters used in the 50% mixtures were retained for consistency, while thresholds were proportionally adjusted for lower purities to account for dilution of unstable reads. Specifically, regions were called unstable when the proportion of deviating reads exceeded ∼half of the simulated tumor fraction (e.g., 5% at 10% tumor purity, 3% at 5% tumor purity, 2% at 3% tumor purity and 1% at 1% tumor purity).

## Data availability

TCGA whole-exome and whole-genome sequencing data used in this study were obtained from TCGA through the GDC and are available to qualified researchers via the GDC and associated dbGaP-controlled access mechanisms in accordance with TCGA data use policies.

Prostate cancer cell-free DNA sequencing data from the PROMIS cohort have been deposited in the European Genome-phenome Archive (EGA) under accession EGAS00001008190 and are available upon data access committee approval. Colorectal cancer cfDNA sequencing data will be made available to qualified researchers upon reasonable request.

Processed MSI scores, locus-level metrics and benchmarking summaries generated by PROMIS are provided as supplementary tables and in the accompanying PROMIS GitHub repository, together with configuration files and example outputs.

## Code availability

The PROMIS pipeline is available at GitHub (https://github.com/vlachosg37/promis). The repository contains the core workflow, configuration templates, curated microsatellite locus sets, example input files, and Conda environment definitions required to reproduce the analyses presented in this study. Documentation in the repository describes supported input data types (whole-exome, whole-genome and targeted panel sequencing), required preprocessing steps, and typical usage scenarios, including cfDNA applications.

## Ethics approval and consent to participate

TCGA data were obtained from The Cancer Genome Atlas (TCGA) via the Genomic Data Commons and used in accordance with TCGA data use policies; all patients provided informed consent as part of the TCGA program.

The prostate cancer cfDNA study was approved by the Ethics Committee of the Medical University of Graz, Austria (approval number: EK Nr ex 21-228). All cfDNA participants provided written informed consent to participate, including collection of blood specimens for plasma genotyping and germline testing and use of associated clinical data for research. The study was conducted in accordance with the Declaration of Helsinki.

## Acknowledgements

We sincerely thank all patients who participated in this study and the clinical teams involved in sample collection and data curation.

## Author contributions

G.V., M.P., J.R.W. and L.A.D. conceived and designed the study. G.V. developed the methodology, performed the formal analysis and investigation, curated resources together with E.H., and carried out validation and visualization with input from T.M. E.H. and L.A.D. secured funding. M.P. and L.A.D. oversaw project administration, and L.A.D. and E.H. provided supervision. G.V. wrote and revised the manuscript.

