## Supplementary for "Reference-free microsatellite instability detection from tumor sequencing using intrasample variability modeling"

### Running title: Tumor-only detection of microsatellite instability

Georgios Vlachos[1,4], Tina Moser[1], Mitesh Patel[4], James R. White[3], Ellen Heitzer[1], and Luis A. Diaz Jr.[4*]

[1]Institute of Human Genetics, Diagnostic & Research Center for Molecular BioMedicine, Medical University of Graz, Austria

[2]Diagnostic and Research Institute of Pathology, Medical University of Graz, Austria

[3]Resphera Biosciences, Baltimore, MD, USA

[4]Division of Solid Tumor Oncology, Memorial Sloan Kettering Cancer Center, New York, NY, USA

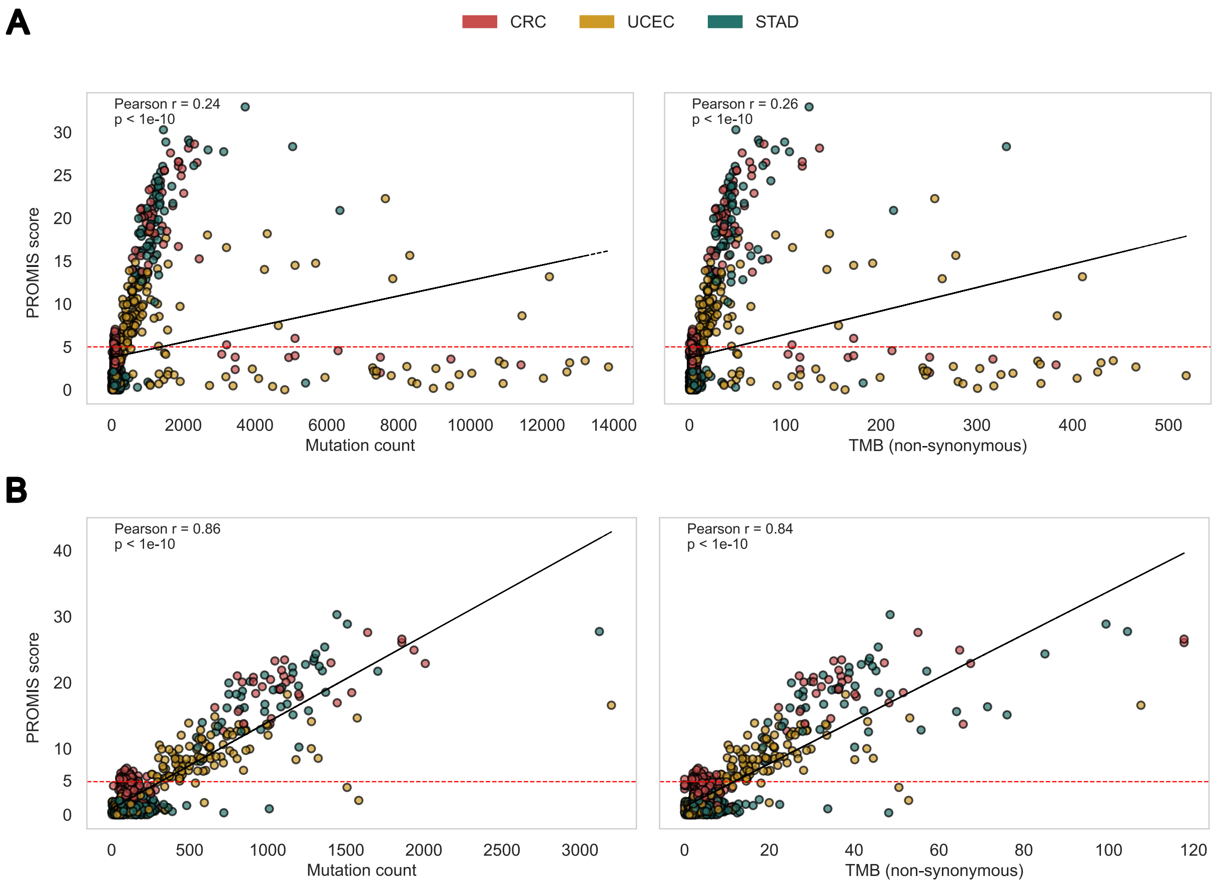


***Supplementary Figure 1. Scatterplots illustrating the relationship between tumor mutational burden and PROMIS score across TCGA colorectal (CRC), endometrial (UCEC), and gastric (STAD) cancers.***

**(A)** Correlation of PROMIS scores with mutation count and nonsynonymous TMB across all cases, including POLE-mutated samples. **(B)** Same analysis after exclusion of POLE-mutated tumors, showing the characteristic reduction of extreme outliers with ultra-high mutational load. Red dashed lines indicate the PROMIS classification threshold (PROMIS = 5).

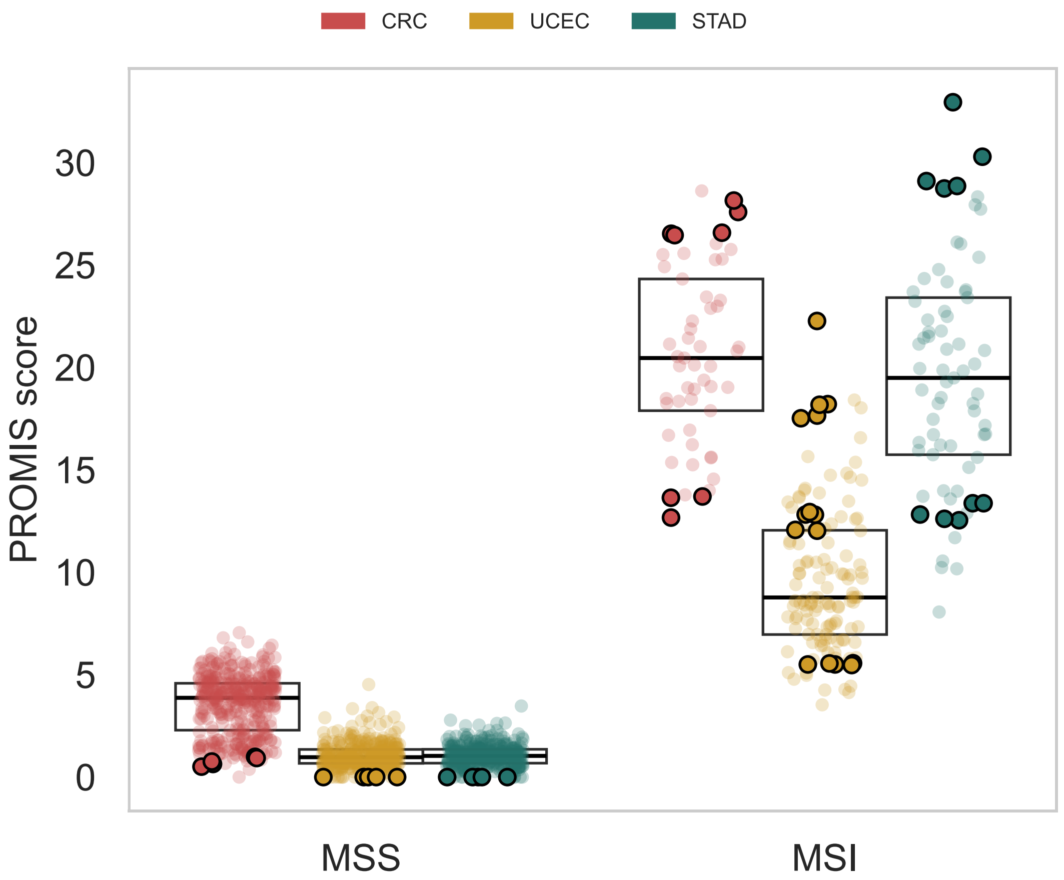


***Supplementary Figure 2. PROMIS score distribution in TCGA colorectal (CRC), endometrial (UCEC), and gastric (STAD) tumors highlighting the 50 representative samples selected for in silico dilution analysis.*** Each point represents a tumor sample, with highlighted points corresponding to the cases used to generate dilution series spanning the full range of PROMIS scores from clearly MSI-H to borderline and MSS classifications. Boxplots indicate the median and interquartile range.

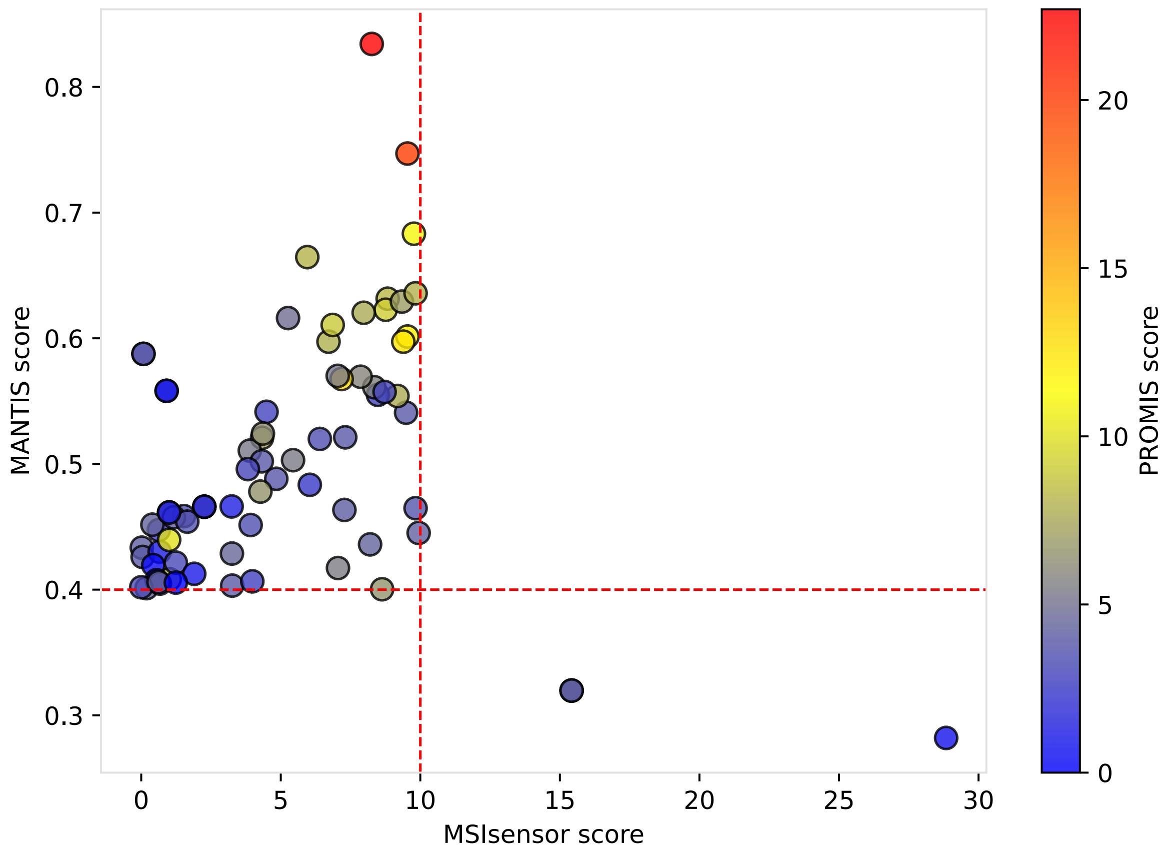


***Supplementary Figure 3. Scatterplot of MSIsensor and MANTIS scores for TCGA colorectal (CRC), endometrial (UCEC), and gastric (STAD) tumors showing cases with discordant classifications between the two conventional MSI callers.*** Each point represents a tumor sample with conflicting MSI status by the established thresholds (MSIsensor > 10 and MANTIS > 0.4; red dashed lines). Points are colored by their PROMIS score using a blue-yellow-red gradient, corresponding to low, intermediate, and high fractions of unstable loci.
